# Single-cell analysis characterizes PLK1 as a catalyst of an immunosuppressive tumor microenvironment in LUAD

**DOI:** 10.1101/2023.08.02.551692

**Authors:** Yifan Kong, Chaohao Li, Jinpeng Liu, Min Zhang, Derek B. Allison, Faisal Hassan, Daheng He, Xinyi Wang, Fengyi Mao, Qiongsi Zhang, Yanquan Zhang, Zhiguo Li, Sai Wu, Chi Wang, Xiaoqi Liu

**Affiliations:** Department of Toxicology and Cancer Biology, University of Kentucky, Lexington, KY, 40536; Department of Biostatistics, University of Kentucky, Lexington, KY, 40536; Markey Cancer Center, University of Kentucky, Lexington, KY, 40536; Department of Pathology and Laboratory Medicine, University of Kentucky, Lexington, KY, 40536

**Author notes:** These authors contribute equally (Yifan Kong, Chaohao Li, Jinpeng Liu). **To whom correspondence should be addressed:** Dr. Xiaoqi Liu: Department of Toxicology and Cancer Biology, University of Kentucky, Lexington, KY, 40536.

**Keywords:** PLK1, LUAD, TME, single-cell RNA-seq, TAM, MHC-II

## Abstract

PLK1 (Polo-like kinase 1) plays a critical role in the progression of lung adenocarcinoma (LUAD). Recent studies have unveiled that targeting PLK1 improves the efficacy of immunotherapy, highlighting its important role in the regulation of tumor immunity. Nevertheless, our understanding of the intricate interplay between PLK1 and the tumor microenvironment (TME) remains incomplete. Here, using genetically engineered mouse model and single-cell RNA-seq analysis, we report that PLK1 promotes an immunosuppressive TME in LUAD, characterized with enhanced M2 polarization of tumor associated macrophages (TAM) and dampened antigen presentation process. Mechanistically, elevated PLK1 coincides with increased secretion of CXCL2 cytokine, which promotes M2 polarization of TAM and diminishes expression of class II major histocompatibility complex (MHC-II) in professional antigen-presenting cells. Furthermore, PLK1 negatively regulates MHC-II expression in cancer cells, which has been shown to be associated with compromised tumor immunity and unfavorable patient outcomes. Taken together, our results reveal PLK1 as a novel modulator of TME in LUAD and provide possible therapeutic interventions.

## Introduction

Lung cancer accounts for one-fifth of all cancer mortalities in the US, making it the leading cause of cancer-related deaths [1]. Lung adenocarcinoma (LUAD) is the major type of this disease, comprising almost half of all cases [2]. LUAD is characterized by a myriad of genetic alternations that drive tumor heterogeneity, such as the common mutation forms involved in receptor tyrosine kinase pathways including EGFR, KRAS, ALK, and ROS1 genes [3]. These mutations exert a profound influence on disease progression and wield considerable sway over patients’ responses to treatment. As a result, a slew of targeted therapies has emerged, tailored to the unique genetic profiles of individual patients. Despite the advances in treatment options, the outlook for LUAD remains dishearteningly bleak, with the 5-year survival rate for metastatic disease is less than 5%. Therefore, precise early screening and the development of more efficacious treatment modalities are essential to improve patients’ outcomes.

Recently, immunotherapies have emerged as a novel treatment option for LUAD [4]. While this marks the new era of cancer treatment, the success of immunotherapies depends on several factors, including the patient’s immune status, and most importantly, the characteristics of the tumor microenvironment (TME), which has emerged as a pivotal area of cancer research [5]. TME is composed of various types of tumor-infiltrating myeloid cells and lymphocytes, and the interactions between cancer cells and these immune cells shape the properties of multiple cancer hallmarks, such as tumor growth, invasion, and metastasis. More closely, immunosuppressive TME can render immunotherapies ineffective, and this nature accounts for the fact that most cancer patients don’t respond to immunotherapy [6]. Within the realm of lung cancer, only 20% of patients respond to monotherapy with checkpoint blockade, a figure that modestly escalates to 40% when paired with conventional chemotherapy [7, 8], underscoring a more complex yet underexplored features of TME in LUAD. Therefore, a better understanding of the TME holds paramount importance in developing effective immunotherapies.

Polo-like kinase 1 (PLK1) emerges as a pivotal serine/threonine kinase, orchestrating a multitude of crucial cellular processes. Its canonical function takes center stage in the intricate regulation of cell division across multiple phases [9–14]. Beyond its canonical role, studies have unveiled its participation in non-mitotic functions, including the regulation of DNA damage repair, metabolism, and immune response [15]. Dysregulation of PLK1, characterized by heightened expression levels, has been identified as a pivotal player in the initiation and progression of various cancer types [16]. Considering its importance in tumorigenesis, novel therapies targeting PLK1 have been developed with observable efficacy in cell and animal models [17]. Both our lab and others have proven that PLK1 overexpression in LUAD leads to cancer progression and is associated with poor overall survival [18, 19]. Interestingly, recent studies have validated PLK1 as a potential target to improve the efficacy of immunotherapies in lung cancer [20], consistent with our finding in pancreatic cancer that targeting PLK1 in pancreatic cancer enhances the effectiveness of immune checkpoint blockade [21]. While these results suggest that PLK1 may be an important modulator of tumor immunity, whether and how PLK1 functions in the TME of LUAD is largely unknown. Here, using single-cell sequencing, we show that elevation of PLK1 incurs a suppressive TME in LUAD, marked by M2 polarization of tumor associated macrophages (TAM) and suppression of antigen presentation process. In addition, elevation of PLK1 attenuates the expression of class II major histocompatibility complex (MHC-II) in professional antigen presentation cells and cancer cells, leading to impaired antigen presentation and anti-tumor immunity. Overall, our results reveal PLK1 as a key regulator of TME and provide evidence of targeting PLK1 to improve immunotherapies in LUAD.

## Materials and Methods

Antibodies, chemicals, and siRNA can be found in **Table S7**.

### Sample preparation for single-cell sequencing

All animal experiments used in this study were approved by the University of Kentucky Division of Laboratory Animal Resources. 8-week-old LSL-Kras^G12D^/Tp53^f/f^ (KP) and LSL-Kras^G12D^/Tp53^f/f^/LSL-Plk1 (KPP) mice were intratracheally instilled with adenovirus-expressing Cre recombinase (Ad-Cre) at a viral titer of 2.5 × 10^7^ PFU per mouse according to the protocol [22]. Tumor formations were monitored every week by MRI and harvested after 12 weeks of adenovirus delivery. Single cell suspensions were prepared following 10x Genomics Cell Preparation Guide. Briefly, mice lung tissues were harvested and cut into smaller pieces, then used pipette to extensively pipette up and down to get tissue mixture in DMEM medium containing 2% FBS. This mixture was filtered with a 70 μm nylon mesh strainer, centrifuged at 300g for 10 mins, then resuspended in DMEM medium containing 2% FBS to get single cell suspension. The single cell suspension was further sorted to enrich immune cell populations, and this suspension was processed for single-cell sequencing. Cell viability was at least 95% for all samples. 10000 cells were targeted for sequencing and loaded onto a Chromium Controller (10x Genomics) for gel beads-in-emulsion formation. Library preparation was conducted using Chromium Next GEM Single Cell 3’ Gene Expression Kit (v3.1, 10x Genomics) according to the manufacturer’s instructions. Single indexed, paired-end libraries were sequenced on an HiSeq 2500 sequencer (Illumina).

### Analysis of single-cell RNA-seq

Single-cell RNA-seq gene expression libraries were mapped to mm10 mouse reference (10x Genomics pre-built reference, mm10-2020-A) using Cell Ranger (v6.1.1, 10x Genomics). For each library, Cell Ranger filtered matrix was subjected to QC filters to remove low quality cells with ≤ 500 features or percentage of mitochondrial transcripts ≥ 15. DoubletFinder analysis was then performed on each library separately to identify and filter potential doublets with default parameters [23]. Additionally, cells with features > 8000 or counts > 50,000 were removed. Pass QC cells from the KP and KPP groups were combined and processed using the Seurat (v3) R package [24]. Count matrices were Log-normalized and scaled. The top 2,000 highly variable genes (HVGs) were selected to define principal components (PCs). Batch integration was performed via Harmony algorithm (v0.1.0) to the combined data for batch effect corrections with default settings [25]. Neighbor analysis was performed by FindNeighbors function using the PCs from the Harmony dimension. Clustering was identified with FindClusters function at resolution = 0.8. UMAP was calculated based on the Harmony dimension for clusters visualization. Identification of cluster markers was performed using a Wilcoxon rank-sum test by comparing each cluster with the rest of the cells. The two-sample binomial test will be used to compare the distribution of cell clusters between the two groups. For differential gene expression analysis, we used DESeq2 test in Seurat [26], to identify the up/down-regulated sets of genes between treatment groups in the cell populations. Gene Set Enrichment Analysis (GSEA) was performed with fgsea R package [27], using KEGG and HALLMARK datasets [28, 29].

### Analysis of public dataset and bulky RNA-seq

For STRING analysis [30], genes functionally associated with CXCL2 were directly checked with online STRING tool (https://string-db.org/). For gene expression, correlation, and survival analyses of TCGA-LUAD dataset, patients’ demographic data with RSEM and zscore values (normalized to all samples) of RSEM were downloaded from cBioportal [31–33]. Analysis of bulky RNA-seq data from KP and KPP tumors was described previously [19].

### Cell culture

KP and KPP cell lines were isolated from transgenic mice 12–14 weeks after adenovirus infection, which was described before [19]. H358 cell line was purchased from ATCC (CRL-5807). All cell lines were cultured in RPMI 1640 medium containing 10% FBS, 1% penicillin-streptomycin at 37°C incubator with 5% CO_2_. For preparation of conditioned medium, KP or KPP cells were cultured in 100mm dishes until 70-80% confluency, then medium was refreshed with DMEM medium containing 0.5% FBS and continued to culture for 48 hours. The resulting conditioned medium was used for coculture experiments and mouse cytokine array. For transfection of siRNA, predesigned siRNA (Sigma) targeting PLK1 were transfected with jetPRIME Versatile DNA/siRNA transfection reagent (Polyplus, 101000001) according to the manufacturer’s instructions. 48 hours after transfection, cells were harvested for subsequent experiments. All the cell lines were within 50 passages and tested negative for mycoplasma contamination.

### Coculture experiments

Macrophages and dendritic cells were isolated from the bone marrows of B6 hosts. Briefly, bone marrows were flushed out of the femur and tibia with complete DMEM medium using 25-gauge needles. After centrifugation at 400g for 5 mins, bone marrows suspension was treated with 1x RBC lysis buffer (Thermo, 00-4333-57) and centrifuged again. The cell pellets were then washed with 1x PBS twice and resuspended in refresh complete DMEM medium. Cells were cultured at 37°C incubator with 5% CO_2_ for a week to induce differentiation. For induction of macrophages, cells were treated with 10ng/ml Csf1. For induction of dendritic cells were treated with 10ng/ml Csf2 and 20ng/ml Il-4. Medium with cytokines was refreshed every two days. For coculture with cells, KP or KPP cells were directly seeded onto the same plates containing immune cells at indicated ratio for 48 hours. Treatment with Cxcl2 or SX-682 started at 6 hours post coculture and continued until end of the experiments. For coculture with conditioned medium, the conditioned medium from KP or KPP, with mouse IgG or anti-Cxcl2 neutralization antibodies, was used to replace normal culture medium for immune cells and continued to culture for 48h.

### Flow cytometry

Cells were harvested by a cell scraper and washed once with 1x PBS, then stained on ice with indicated antibodies (1:1000) in 1x PBS for 30 mins. Samples containing tumor cells were fixed with 70% ethanol before staining. Data were acquired on the BD Symphony A3 analyzer (BD Biosciences) and analyzed using FlowJo software (BD Biosciences).

### Mouse cytokines array

Mouse cytokine array experiment was performed using Proteome Profiler Mouse Cytokine Array Kit (R&D Systems, ARY006) according to the manufacturer’s instructions. Briefly, conditioned medium from KP and KPP was incubated with membranes containing pre-adsorbed primary antibodies targeting mouse cytokines. Following wash step, the membranes were then incubated with HRP-linked secondary antibodies. After washing membranes again, membranes were probed with chemiluminescent reagents to visualize cytokine spots. Medium containing 0.5% FBS but without cancer cells was used as negative control. Images were captured with ChemiDoc Imaging System (Bio-Rad) and analyzed with Image Lab software (Bio-Rad).

### Immunohistochemistry (IHC)

Formalin-fixed, paraffin-embedded slides prepared from KP and KPP tumors were stained with MHC-II antibodies. IHC staining was performed with the VECTASTAIN Elite ABC Universal PLUS Kit (Vector Laboratories, PK-8200) according to the manufacturer’s instructions. Staining was visualized with ImmPACT DAB Substrate Kit (Vector Laboratories, SK-4105) and counterstained with Harris’s hematoxylin. Images were with the Nikon microscope.

### Statistical analysis

Statistical analyses were performed with the statistical functions in GraphPad Prism 8 and R programming language. Unless denoted elsewhere, an unpaired two-sided t test was used as the default method for results. Normality and variance of results were checked to confirm the requirements of t test. Statistical significance was set at p < 0.05 unless denoted otherwise.

## Results

### Elevation of PLK1 is associated with immunosuppressive TME in LUAD

To investigate the impact of PLK1 on lung cancer TME, we designed a workflow to perform the single-cell RNA-seq using our LSL-Kras^G12D^/Tp53^f/f^ (KP) and LSL-Kras^G12D^/Tp53^f/f^/LSL-Plk1 (KPP) mouse models (**Fig. 1A**), which were previously reported [19]. Using the markers to differentiate non-immune and immune cells (**Figs. S1A-S1B**), the results showed that most cells (over 95%) were captured as immune cells without significant differences in KP and KPP groups (**Fig. S1C**), demonstrating the efficacy and reliability of single-cell data to characterize TME in both groups. Unsupervised clustering of immune cells generated 20 different immune cell clusters in samples characterized by a specific gene signature (**Figs. S1D-S1E, Table S1**), indicating a significant intratumoral heterogeneity of immune cell populations. Notably, the different proportions of immune cell clusters suggested a distinct TME in KP and KPP (**Figs. S1F-S1G**). To better visualize the differences in immune cell populations, we performed a supervised clustering of major immune cell types based on their specific gene signatures (**Figs. 1B-1C, Table S2**). Results showed that KPP tumors had significantly lower proportions of all cell types except for TAM compared to KP tumors (**Figs. 1D-1E**). In both groups, TAM were the major population, suggesting the unique function of them in lung cancer progression. Besides, the lower levels of tumor-infiltrating lymphocytes (B cells and T cells) in KPP tumors indicated an immunosuppressive TME in this group. Since T cells and NK cells are the major immune cells exerting direct anti-tumor immunity, we further investigated these populations by clustering them in KP and KPP tumors (**Figs. S2A-S2B, Table S3**). Clearly, KPP tumors displayed decreased proportions of tumor-infiltrating NK cells and T cells of all subtypes (**Fig. S2C**). The lower levels of these tumor-infiltrating lymphocytes in KPP tumors were impactful and further supported a cold TME in this group, as they were the major immune cells responsible for eliminating tumors. This observation suggested that KPP tumors were specifically protected from the anti-tumor immunity and thus resistant to the host immune defense. Taken together, these results illustrated that high PLK1 in LUAD promoted a suppressive immunophenotype.

**Figure 1.**
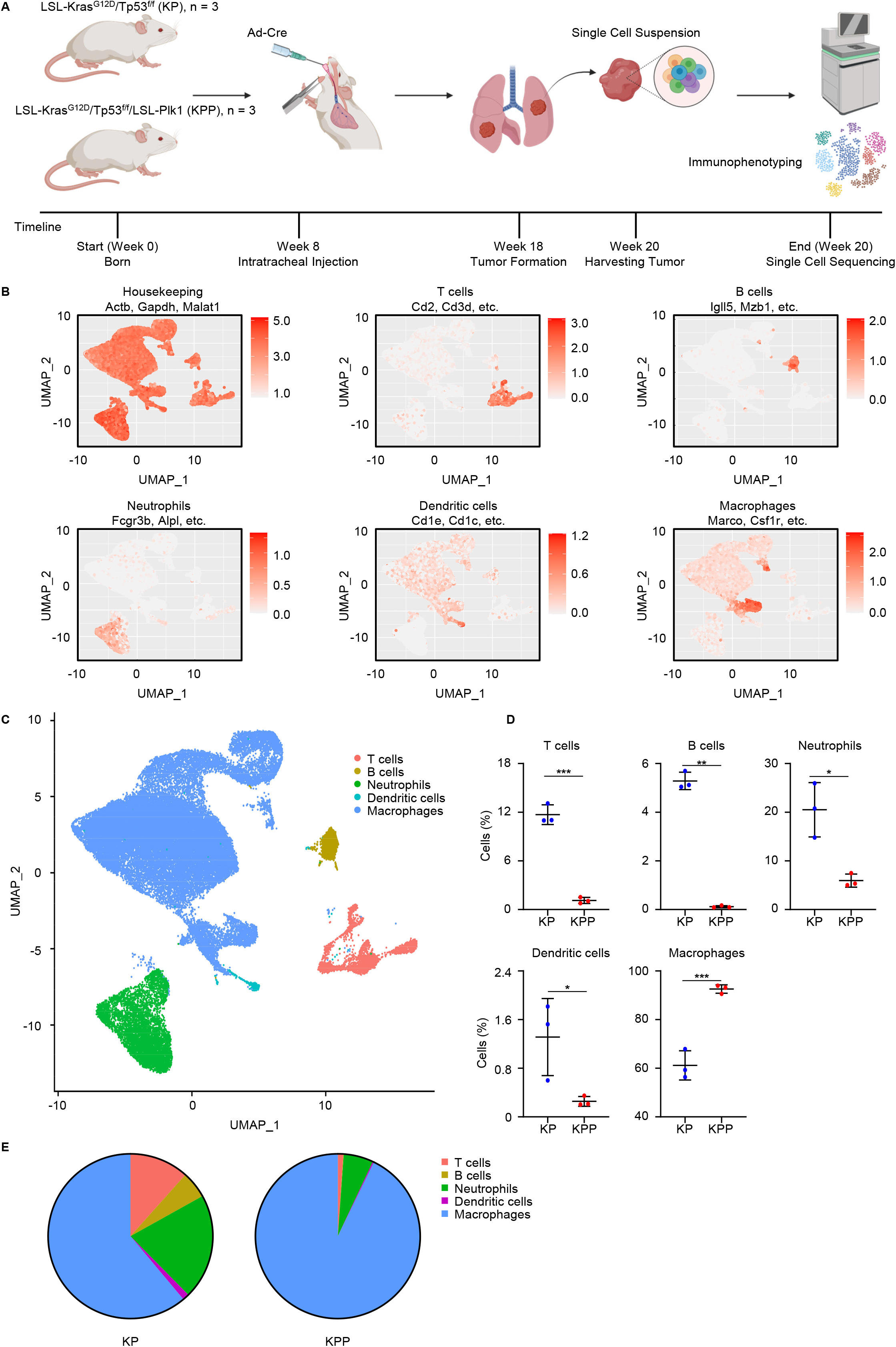
Single-cell RNA-seq analysis of KP and KPP mice. **A,** Workflow of single-cell RNA-seq of KP and KPP mice. **B,** Representative genes used for classification of T cells, B cells, Neutrophils, Dendritic cells, Macrophages. **C,** UMAP of major immune cell populations in KP and KPP. **D,** Comparison of cell populations between KP and KPP (n=3). **E,** Proportion of cell populations in KP and KPP. *, p < 0.05. **, p < 0.01. ***, p < 0.001.

### Tumor-promoting M2 macrophages propagate in high-PLK1 TME of LUAD

The relatively higher proportion of TAM in KPP tumors was intriguing, and this phenomenon enticed us to investigate the role of TAM in high-PLK1 TME. Given that the functions of TAM were dichotomous, we specially focused on the proportions of M1 and M2 macrophages in KP and KPP tumors, which were tumor-suppressing and tumor-promoting, respectively. Analysis of gene expressions in TAM identified 1201 differentially expressed genes with statistical significance between KP and KPP tumors (**Fig. 2A, Table S4**). Investigation of top 10 up and down regulated genes in KPP group revealed that TAM in KPP tumors were positively enriched for genes associated with M2 macrophages (e.g., Chil3, Ccn3) while negatively enriched for genes associated with M1 macrophages (e.g., H2-Eb1, H2-Ab1), suggesting that TAM in KPP tumors were more enriched for M2 subtype (**Fig. 2B**). Comparison of M1 gene signature and M2 gene signature in KP and KPP groups further supported increased M2 macrophages in KPP tumors (**Fig. 2C, Table S5**). To confirm this phenotype, we performed supervised clustering of total macrophages (**Figs. 2D-2E, Table S5**), and the results indeed demonstrated that TAM in KPP tumors consisted of more M2 macrophages compared to KP tumors (**Figs. 2F**). Since M2 macrophages were considered tumor-promoting and immunosuppressive, the higher proportion of M2 macrophages in KPP tumors was consistent with a colder TME in this group. In sum, these results supported the notion that high PLK1 was associated with increased tumor-promoting M2 macrophages in LUAD.

**Figure 2.**
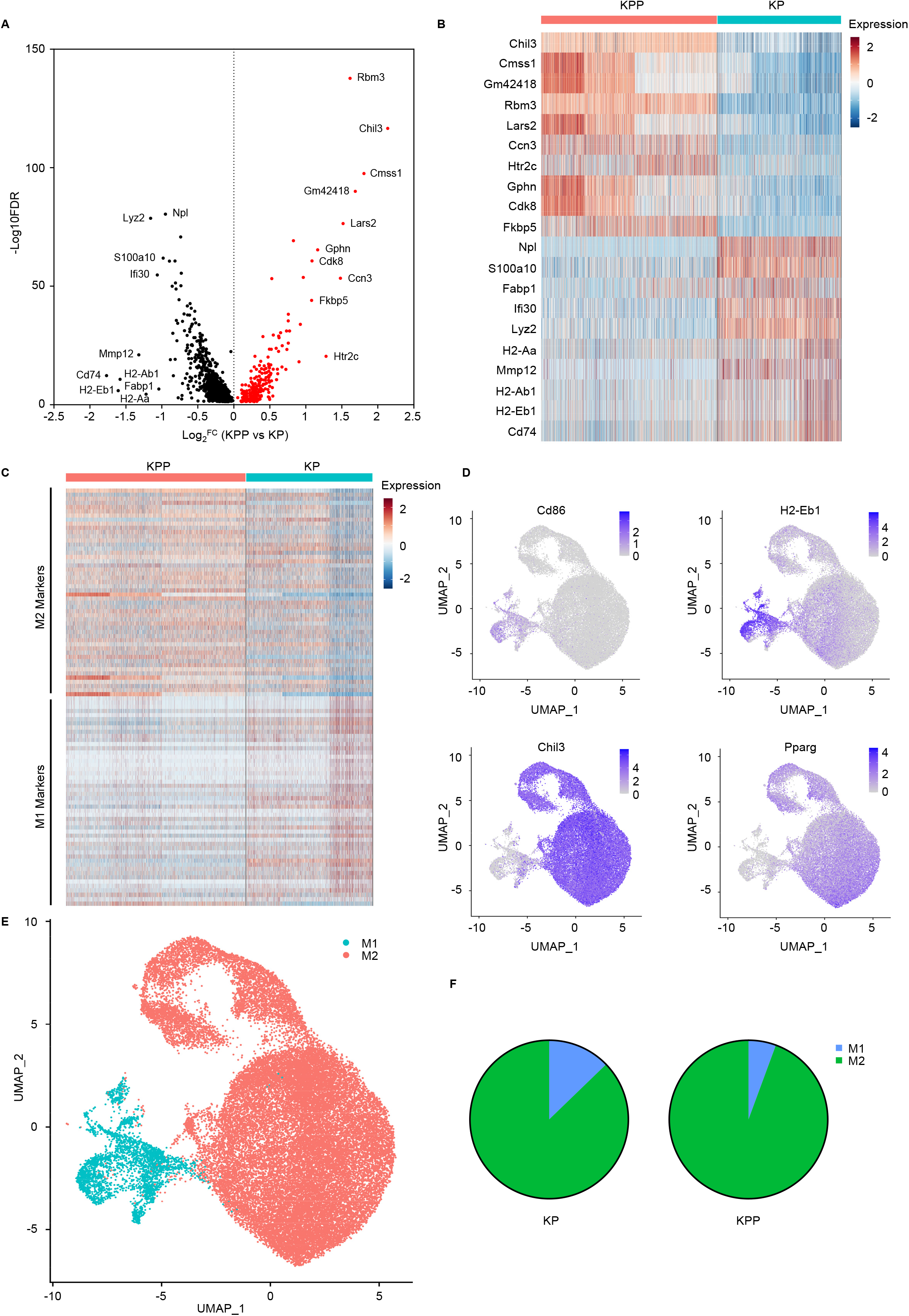
High PLK1 is associated with increased M2 macrophages. **A,** Volcano plot of differentially expressed genes (KPP vs KP, FDR < 0.05) in TAM. Top 10 up and down regulated genes (ranked by Log_2_^FC^) are labeled. Log_2_^FC^, Log_2_^fold change^. **B,** Heatmap of top 10 up and down regulated genes identified in A. **C,** Heatmap of top 50 representative genes (ranked by Log_2_^FC^) in M1 and M2 gene signatures. **D,** Representative genes and functional markers used for classification of M1 (Cd86, H2-Eb1) and M2 (Chil3, Pparg) macrophages. **E,** UMAP of M1 and M2 macrophages in KP and KPP. **F,** Proportion of M1 and M2 macrophages in KP and KPP.

### High PLK1 suppresses antigen presentation pathway and induces M2 polarization

Since high PLK1 was associated with increased infiltration and differentiation of M2 macrophages, we sought to delineate the biological consequence of this phenotype. To achieve this aim, we performed GSEA in TAM. The results showed that multiple pathways associated with the functions of macrophages were dramatically disturbed (**Fig. 3A, Table S6**). Several pathways related to immune response were suppressed in KPP tumors, such as pathways associated with inflammation (e.g., IFN-γ response, **Fig. 3B**), pathogen defense (e.g., viral myocarditis), and autoimmune diseases (e.g., systemic lupus erythematosus). In addition, pathways associated with steroid and lipid metabolism were elevated in KPP tumors (e.g., steroid biosynthesis, **Fig. 3B**), which were previously demonstrated to promote M2 polarization [34]. These results suggested an impaired anti-tumor immune defense and were in congruent with the observations of more M2 macrophages and a more immunosuppressive phenotype in KPP tumors. Of note, antigen processing and presentation pathway was among the most suppressed pathway in TAM (**Figs. 3A-3B, Table S6**). Professional antigen presentation cells such as macrophages rely on MHC-II and costimulatory factors for their antigen processing and presentation function. Indeed, review of gene expressions and macrophages markers (**Figs. 2A-2C, Table S4, Table S5**) showcased that some low-expression M1 genes in TAM of KPP tumors were also MHC-II genes (e.g., Cd74, H2-Eb1, H2-Ab1, H2-Aa) and costimulatory factors (e.g., Cd86, Cd80), consistent with the impaired antigen presentation function in TAM. Given that this pathway was key to successful anti-tumor immunity [35], the loss of antigen presentation function in TAM and more tumor-promoting M2 macrophages in KPP group supported the immunosuppressive role of PLK1. Moreover, results of GSEA in other immune cells showed that dampened antigen processing and presentation was a shared consequence among immune cell populations (**Fig. S3, Table S6**), suggesting that suppression of this critical pathway was a general consequence of high PLK1 expression. Based on these results, we hypothesized that PLK1 could inhibit antigen presentation pathway, and this led to M2 polarization. To validate our hypothesis, we performed in vitro coculture experiments using established cell lines from KP and KPP mice and used flow cytometry to monitor expressions of important genes and markers in antigen presentation and macrophages polarization **(Figs. 3C-3D)**. Short-term coculture of bone marrow derived macrophages (BMDM) with KP or KPP showed that both cells elevated costimulatory factors (Cd80 and Cd86) with even stronger effect from KPP group. However, only KPP cells were able to induce higher level of M2 marker Cd206, indicating the M2 polarization effect of PLK1. Coculture with both cell lines attenuated MHC-II expression, possibly due to the immune suppression mechanism of tumors. Compared to KP, the level of MHC-II was lower in KPP group. Given that MHC-II plays a pivotal role in antigen processing and presentation, the lower expression of MHC-II in KPP group confirmed that antigen presentation function was specifically dysregulated in high PLK1 condition. Collectively, these data demonstrated that PLK1 suppressed antigen presentation pathway and induced M2 polarization in LUAD.

**Figure 3.**
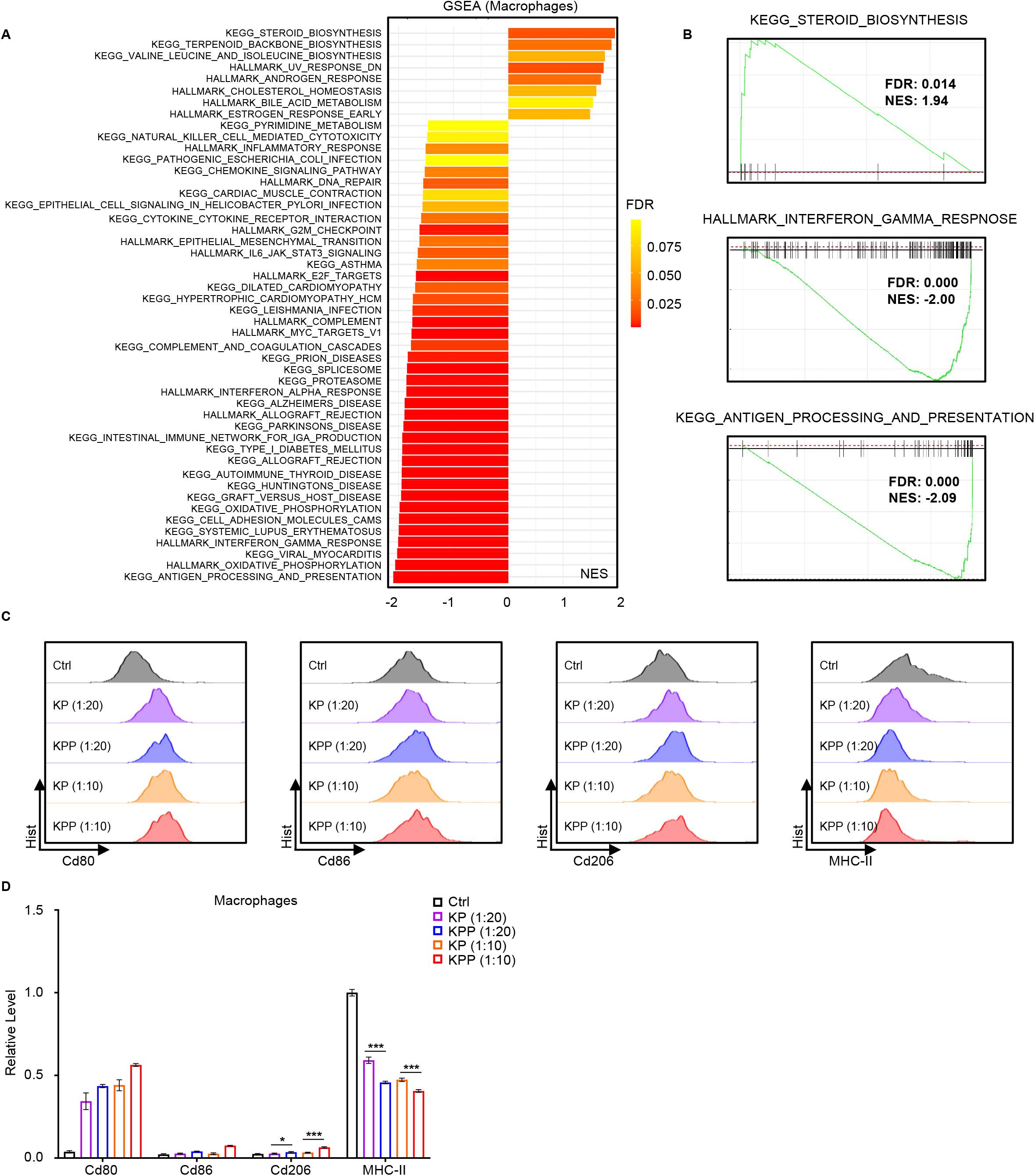
PLK1 suppresses antigen presentation pathway and promotes M2 polarization. **A,** Significant pathways (KPP vs KP, FDR < 0.1) altered in macrophages identified by GSEA. NES, normalized enrichment score. **B,** GSEA plots of three exemplary pathways in A. **C,** Flow cytometry analysis of indicated genes in BMDM cocultured with KP or KPP at the indicated ratio for 48 hours. **D,** Quantification of C (n=3). *, p < 0.05. ***, p < 0.001.

### Immunosuppressive function of PLK1 depends on CXCL2

Paracrine signaling through the release of cytokines from cancer cells is critical for intercellular communication and shaping anti-tumor immunity [36]. Inspection of pathway analysis in TAM (**Fig. 3A, Table S6**) revealed that several cytokine and chemokine pathways were altered (e.g., interferon response pathway), and this led to our hypothesis that PLK1 exerted its immunosuppressive function via regulation of cytokine secretion. To test our hypothesis, we performed a cytokines array of mouse cytokines released from the conditioned medium of KP and KPP cells. We found that cytokine profiles were indeed different between KP and KPP. Compared to KP, KPP cells secreted more Il-6, Cxcl2 and Ccl3 cytokines, but less Ccl5 (**Fig. 4A**). The increased secretion of Il-6 was reasonable given its established role in M2 polarization [37]. Besides, Ccl3 (macrophage inflammatory protein 1-alpha) was one of the cytokines released from macrophages and Ccl5 was responsible for T cells chemotaxis [38, 39], so their alternations were congruent with the higher proportion of macrophages and lower infiltration of T cells in KPP tumors. Among them, we were especially interested in Cxcl2, as it was the most highly secreted cytokine in the supernatant of KPP. It has been shown that secreted CXCL2 in TME can recruit myeloid derived suppressor cells and M2 macrophages [40, 41], and this leads to suppression anti-tumor response (**Fig. 4B**). We also investigated which pathway was responsible for the increased secretion of Cxcl2 in KPP. A preliminary STRING analysis in human and mice identified several important genes related to CXCL2 (**Figs. S4A**). Except for some known interaction genes (e.g., CXCR1 and CXCR2, two receptors for CXCL2), we found that in both species the genes related to JAK-STAT3 pathway (e.g., IL6/Il6) and NFKB pathway (e.g., TNF, Rela) commonly emerged. GSEA of RNA-seq data confirmed that two pathways were highly active in KPP cells (**Fig. S4B**). Of note, although not very striking, Il-6 was simultaneously detected with Cxcl2 in the secretion profile of KPP, which was not observed in KP control (**Fig. 4A**). These results provided evidence that IL-6/JAK/STAT3 and TNF/NFKB axis might account for the increased Cxcl2 in KPP cells.

**Figure 4.**
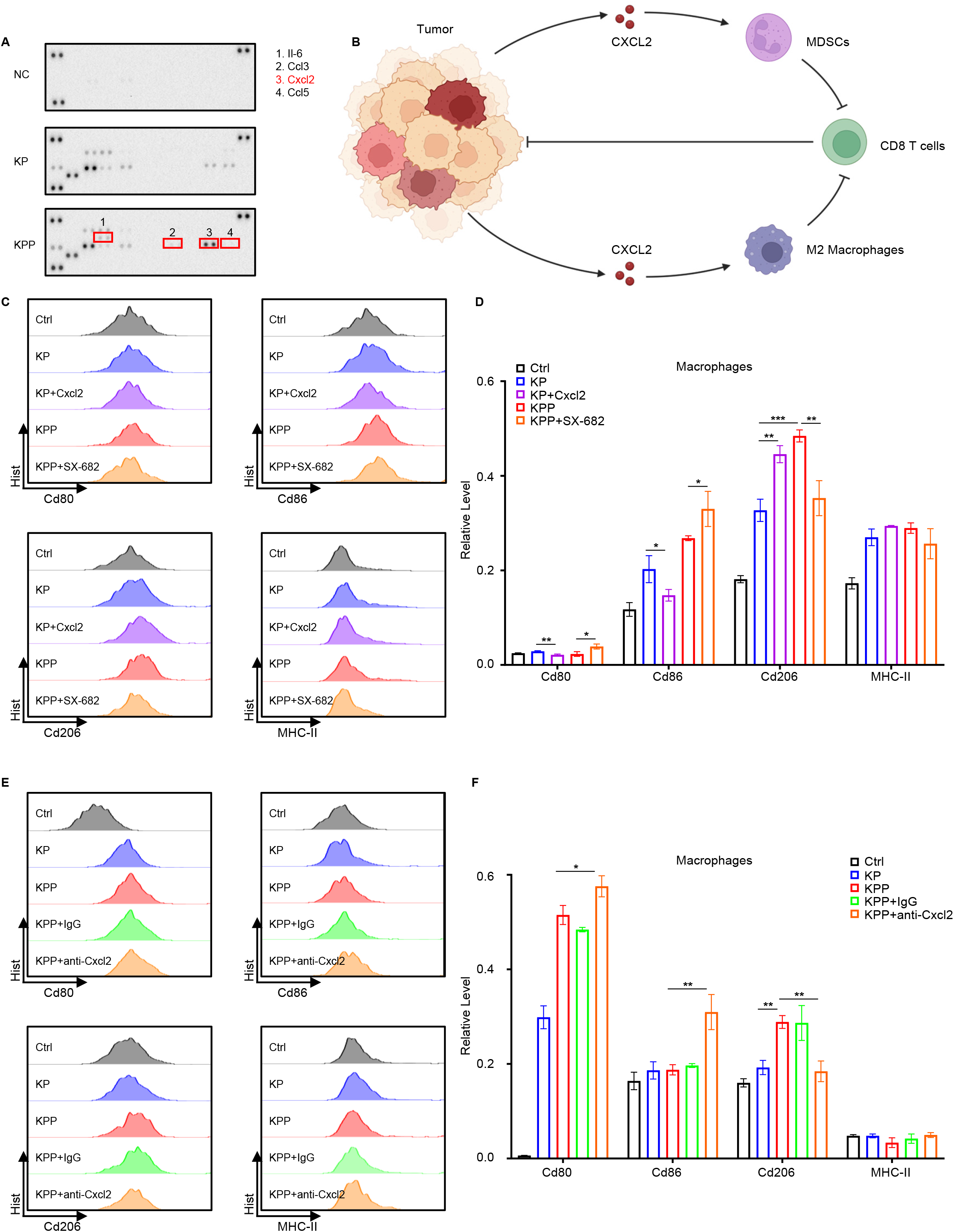
CXCL2 is responsible for the immunosuppressive function of PLK1. **A,** Cytokines array detection of cytokines secretion from the culture medium of KP and KPP. Notable differences are labeled. **B,** Illustration of CXCL2’s function in suppressing anti-tumor immunity. **C,** Flow cytometry analysis of BMDM cocultured with conditioned medium from KP or KPP for 48 hours, with or without recombinant Cxcl2 (2ug/ml) or CXCR1/CXCR2 inhibitor SX-682 (1µM) treatment. **D,** Quantification of C (n=3). **E,** Flow cytometry analysis of BMDM cocultured with conditioned medium from KP or KPP for 48 hours, with or without anti-Cxcl2 neutralization antibodies (5ug/ml) treatment. **F,** Quantification of E (n=3). *, p < 0.05. **, p < 0.01. ***, p < 0.001.

In mice, Cxcl2 can bind to Cxcr1 and Cxcr2, albeit with much higher affinity to Cxcr2 [42]. Given that Cxcl2 was predominant in KPP-secreted cytokines and its established role in recruiting M2 macrophages, we hypothesized that the observed immunosuppressive function of PLK1 depended on CXCL2. To test this assumption, we performed coculture experiments of BMDM with conditioned medium from KP or KPP (Figs. 4C-4D), treated with recombinant mouse Cxcl2 or SX-682, a previously reported dual CXCR1/CXCR2 inhibitor in clinical trial [43]. Coculture of BMDM with conditioned medium recapitulated most of the results from coculturing with cells, but MHC-II was not decreased. Instead, MHC-II elevated under this condition, indicating that the decrease of MHC-II in coculture with cells directly came from interaction between cancer cells and immune cells. Coculture with conditioned medium still elevated Cd80 and Cd86 in both groups, with a much higher induction of M2 marker Cd206 in KPP group. Intriguingly, the addition of Cxcl2 in conditioned medium from KP was able to reduce Cd80 and Cd86 while increasing the level of Cd206. Besides, adding CXCR1/CXCR2 inhibitor in conditioned medium from KPP group elevated Cd80 and Cd86 but attenuated Cd206 expression. However, MHC-II was not disturbed by either Cxcl2 or SX-682. Considering that mouse Cxcl2 binds to Cxcr2 with much higher affinity, and mouse Cxcr1 is mainly for Cxcl6 [42], the off-target effect of SX-682 in our experimental setting could be ignored. To address this issue, we performed a parallel coculture experiment of BMDM with conditioned medium from KP and KPP (**Figs. 4E-4F**), treated with mouse IgG control or anti-Cxcl2 neutralization antibodies. Treatment with neutralization antibodies witnessed an uptrend of Cd80 and Cd86 but a downtrend of Cd206, with an unchanged status of MHC-II, reassuring the results obtained from the coculture experiment using SX-682. These data illustrated that Cxcl2 was partially responsible for the effect of PLK1 in suppressing antigen presentation and inducing M2 polarization, mainly focused on regulation of costimulatory factors. As costimulatory factors in professional antigen presentation cells, the functions of CD80 and CD86 were similar in dendritic cells, and CD206 was also a hallmark of immature dendritic cells with incomplete immune functions [44]. Thus, detection of these markers in dendritic cells has similar value in dissecting the biological effect of PLK1 and CXCL2 on antigen presentation. Thus, we repeated all three coculture experiments with dendritic cells isolated from bone marrow. The results revealed that alternations of Cd80, Cd86 and Cd206 were quite similar and repeatable, but the level of MHC-II displayed distinct changes. In the coculture experiment with KP/KPP cells and dendritic cells, an increased proportion of KP cells accompanied an elevated level of MHC-II, but an increased ratio of KPP cells reduced MHC-II expression (**Figs. S5A-S5B**). This was different from the results in BMDM, where a higher ratio of both cell lines was associated with a lower level of MHC-II. However, in both settings, KPP cells were capable of diminishing MHC-II expression more strongly compared to KP, suggesting PLK1’s universal suppression of antigen presentation by limiting MHC-II expression. In the coculture experiment with KP/KPP conditioned medium and dendritic cells, the results demonstrated that the effect of PLK1 on MHC-II could be explained by CXCL2, as treatment with Cxcl2 in KP group reduced MHC-II expression and treatment with either SX-682 or neutralization antibodies in KPP group elevated MHC-II level (**Figs. S5C-S5F**), which was different from the results in macrophages. Despite these differences, the disturbance of antigen presentation and induction of immunosuppressive environment by CXCL2 was observed. Taken together, these results stated that CXCL2 accounted for the immunosuppressive function of PLK1 in LUAD, but the actual effect of CXCL2 might vary in different immune cells.

### PLK1 negatively regulates MHC-II in cancer cells

It has been reported that MHC-II can also express on the surface of cancer cells, and this phenotype is associated with improved response to immunotherapy and better survival of cancer patients [45–48]. However, this phenotype was unexplored in lung cancer. Based on these established facts and our observations that PLK1 was negatively associated with MHC-II, we aimed to explore whether PLK1 affected the expression of MHC-II in LUAD tumors. We first detected MHC-II expression by IHC staining on tumor slides prepared from KP and KPP mice, and the results indeed showed that both tumors expressed MHC-II, with more striking expression in KP tumors (**Fig. 5A**). This was consistent with the results observed in immune cells that high PLK1 led to lower MHC-II level. Next, we performed flow cytometry analysis of MHC-II level in KP and KPP cell lines, and the results verified the lower expression of MHC-II in KPP cells (**Figs. 5B-5C**). Since MHC-II expression in cancer cells depends on the IFN-γ [49], we performed ELISA experiment to detect intracellular IFN-γ levels between KP and KPP cells. As expected, KPP cells had a lower level of IFN-γ compared to KP (**Fig. 5D**), consistent with its inferior expression of MHC-II. To direct assess the effect of PLK1, we treated KP and KPP with three PLK1 inhibitors (PLKi), including BI-2536 (BI), GSK461364A (GSK) and Onvansertib (ONV). In both cells, treatment with PLK1i consistently elevated the level of MHC-II (**Figs. 5E-5H**), confirming the negative regulation of MHC-II by PLK1. To assess the human side, we utilized human H358 LUAD cell line to see whether PLK1 played a similar role. We first transfected H358 cells with siRNA targeting PLK1, and the depletion of PLK1 elevated MHC-II as expected (**Figs. 5I-5J**). Dose escalation of PLKi also showed a gradual increase in MHC-II expression (**Figs. 5K-5L**), reassuring the effect of PLK1. Cumulatively, these data supported the conclusion that PLK1 also regulated MHC-II in LUAD cancer cells.

**Figure 5.**
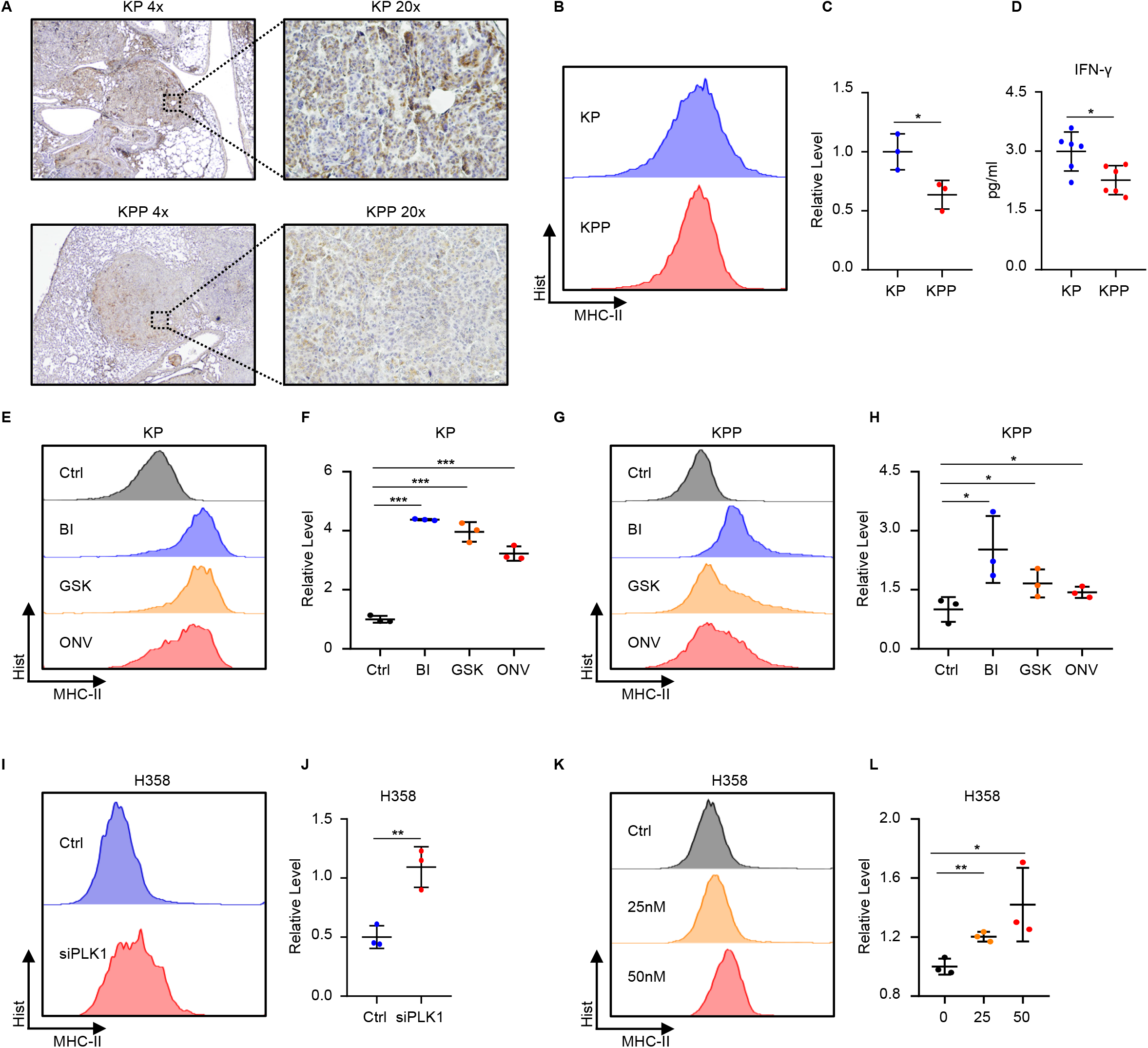
PLK1 suppresses MHC-II expression in tumor cells. **A,** Representative IHC images of KP and KPP tumors stained with MHC-II antibodies. **B,** Flow cytometry detection of MHC-II in KP and KPP cells. **C,** Quantification of B (n=3). Data are normalized to KP and shown as mean±SD. **D,** ELISA detection of IFN-γ in KP and KPP cells (n=6). Data are normalized to KP and shown as mean±SD. **E,** Flow cytometry detection of MHC-II in KP cells treated with PLK1 inhibitors (50nM BI, 50nM GSK, 200nM ONV) for 48 hours. **F,** Quantification of E (n=3). Data are normalized to Ctrl and shown as mean±SD. **G,** Flow cytometry detection of MHC-II in KPP cells treated with PLK1 inhibitors (50nM BI, 50nM GSK, 200nM ONV) for 48 hours. **H,** Quantification of G (n=3). Data are normalized to Ctrl and shown as mean±SD. For these results, an unpaired one-sided t test was used. **I,** Flow cytometry detection of MHC-II in H358 cells 48 hours after transfection with siRNA control or siRNA targeting PLK1. **J,** Quantification of I (n=3). Data are normalized to Ctrl and shown as mean±SD. **K,** Flow cytometry detection of MHC-II in H358 cells with dose escalation of PLK1 inhibitor ONV (nM) for 48 hours. **L,** Quantification of K (n=3). Data are normalized to untreated group (0) and shown as mean±SD. *, p < 0.05. **, p < 0.01. ***, p < 0.001.

### Clinical assessment of PLK1’s regulation of MHC-II

We’ve previously demonstrated that high PLK1 was associated with worse patients’ outcomes [19], and this was consistent with the data that PLK1 suppressing MHC-II presented in this study. To evaluate the translational values of our findings, we analyzed the TCGA-LUAD dataset. We found that high PLK1 was associated with low levels of MHC-II member genes (**Figs. 6A, S6**).

**Figure 6.**
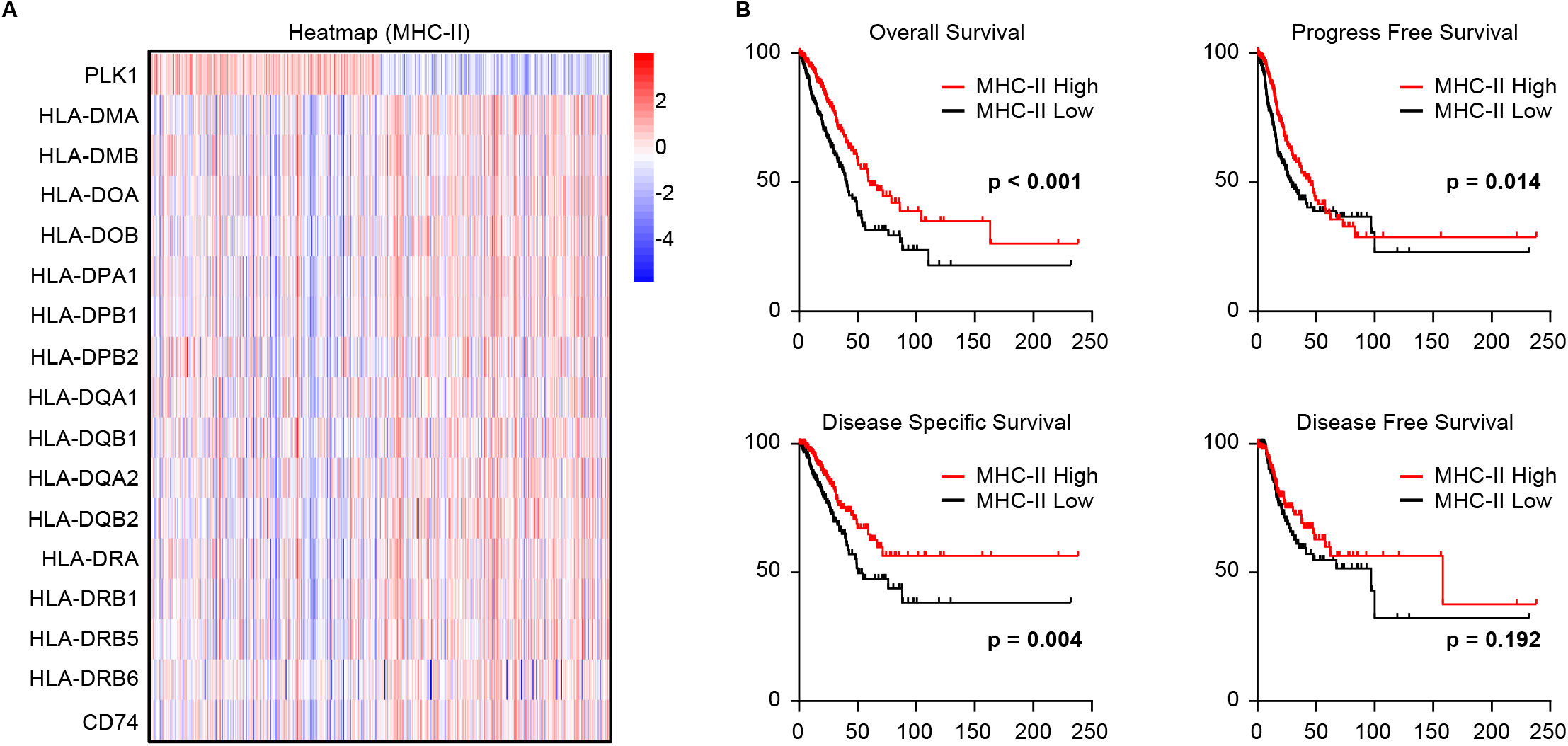
Clinical evidence of PLK1 in suppressing MHC-II in lung cancer. **A,** Heatmap of MHC-II signature genes in TCGA-LUAD patients separated into high PLK1 and low PLK1 groups. Patients are clustered based on the median expression (zsocres of Log_2_^RSEM+1^) of PLK1. **B,** Survival curves of TCGA-LUAD patients separated into MHC-II high and MHC-II low groups. Log_2_^RSEM+1^ of MHC-II signature genes in A are averaged to get signature scores. Patients are clustered based on the median value of signature scores. For these results, a Log-rank test was used.

Using this MHC-II gene signature, we calculated the MHC-II signature scores of all patients and used these signature scores to re-classify patients into high MHC-II and low MHC-II groups, then performed survival analysis of those patients. We found that patients with high MHC-II scores had better survival outcomes compared to patients with low MHC-II scores (**Fig. 6B**). These results were consistent with our previous findings and supported the notion that the negative regulation of MHC-II by PLK1 may promote LUAD progression and result in poor outcomes. In conclusion, PLK1 was negatively associated with MHC-II in clinical situations, and this had an impact on patients’ survival possibly through suppressing antigen presentation pathway.

## Discussion

Here in this study, we render pioneering data to support the fact that PLK1 promoted an immunosuppressive TME in LUAD (**Fig. 7**). Under low PLK1 condition, TME is characterized by a higher proportion of tumor infiltrating lymphocytes and lower level of CXCL2, and this condition is associated with more M1 polarization and functional antigen presentation pathway. In addition, tumors express more MHC-II and this feature is associated with better patient’s survival. Under high PLK1 condition, TME is characterized by low tumor infiltrating lymphocytes and high secreted CXCL2, which promotes M2 polarization and disrupts antigen processing and presentation. Tumors in this situation also express a lower level of MHC-II, which is associated with worse outcomes of patients. The differences in TME shown in high PLK1 and low PLK1 conditions suggest that PLK1 is an important modulator of TME, and targeting PLK1 may be a practical therapeutic intervention in LUAD treatment.

**Figure 7.**
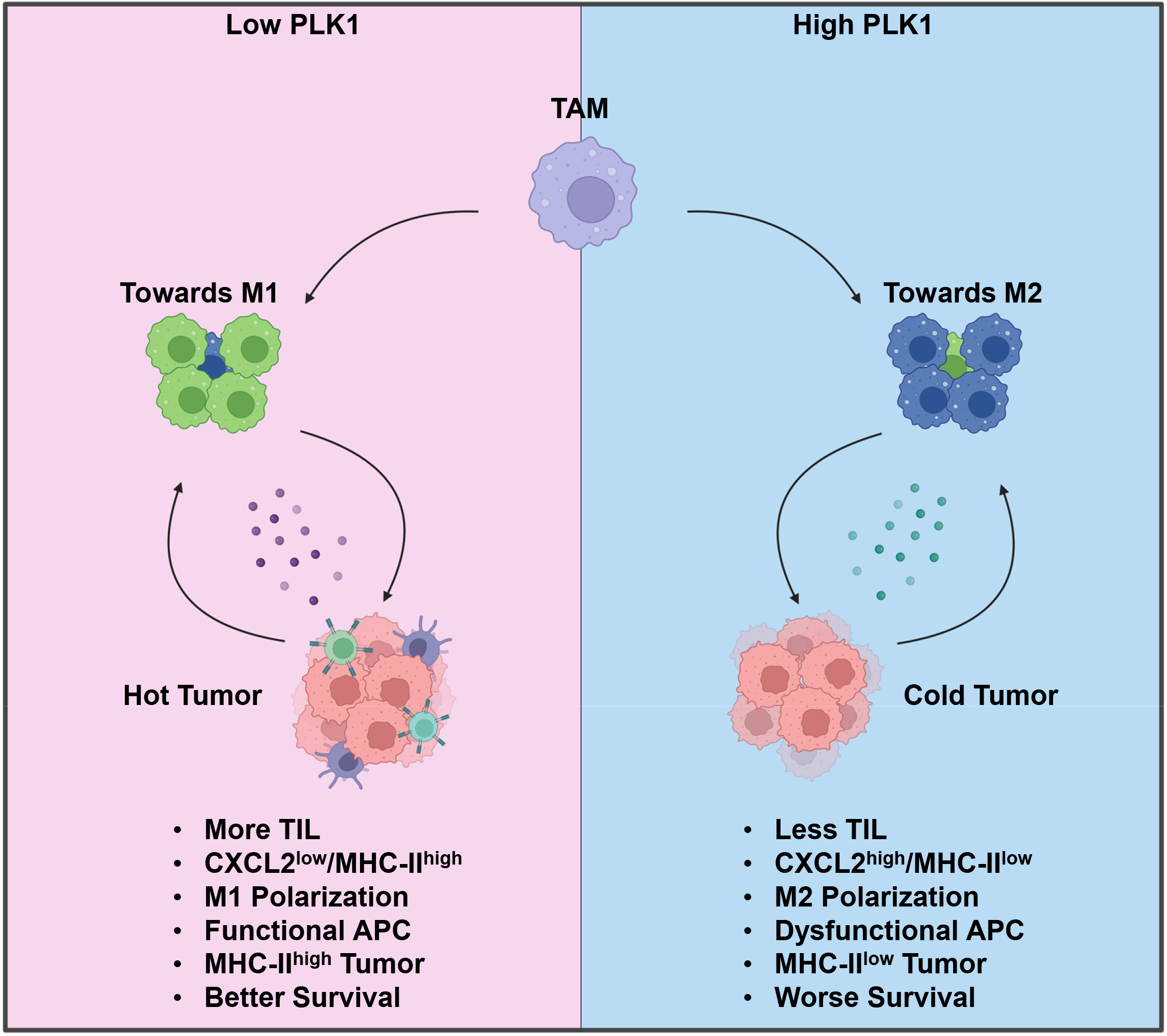
Summary of working model. Under low PLK1 condition, there is low level of CXCL2 secreted from tumors. Tumor microenvironment is “hot”, characterized by increased tumor infiltrating lymphocytes, M1 polarization of TAM and functional APC. Cancer cells are MHC-II high and this feature is associated with better survival of patients. Under high PLK1 condition, there is increased secretion of CXCL2 from tumors. Tumor microenvironment is “cold”, characterized by low levels of tumor infiltrating lymphocytes, M2 polarization of TAM and dysfunctional APC. Cancer cells are MHC-II low and patients have worse survival outcomes.

Although the emergence of immunotherapy has expanded the treatment options for lung cancer patients, the efficacy is questionable due resistance to immunotherapy. Thus, identifying mechanisms behind this harmful condition is urgently needed. Here, we report a novel mechanism of PLK1 in cancer promotion. Previously, PLK1 has been largely considered as a canonical cell cycle regulator in the past decades. However, recent studies have gradually uncovered the tumor-promoting functions of this kinase. The fact that PLK1 suppresses anti-tumor in LUAD is intriguing given that immunotherapy is one of the major therapeutic approaches for lung cancer. The major reasons of resistance to immunotherapies are the scarcity of tumor-infiltrating immune cells and existence of immune checkpoints [50]. We show that high PLK1 is associated with lower infiltrating cytotoxic T cells and NK cells, and this feature is harmful to anti-tumor response in LUAD. The decreased infiltration of T cells and NK cells may be due to the increased M2 polarization from TAM, which is a well-established suppressor of their functions. Besides, functional antigen presentation in TME is the key to the success of anti-tumor immune response, and immune evasion through suppressing this critical process significantly reduces the efficacy of immunotherapy [35]. The novelty of our study lies in the fact we first demonstrate that PLK1 inhibits antigen presentation pathway by down regulating MHC-II in professional antigen-presenting cells, which is the key molecule responsible for processing and presentation of tumor-specific neoantigens. Besides, this conclusion can be extended to cancer cells, where high PLK1 is also associated with low MHC-II expression. The expression of MHC-II in tumors is a good biomarker for better survival of patients [45–48]. However, the function of MHC-II in cancer cells is unclear. Given that MHC-II is mainly responsible for antigen presentation, the fact that MHC-II expressing in cancer cells may be involved in this process by functionally mimicking antigen presentation process and priming CD4 T cells, which directly bind to MHC-II to activate immune response, and this idea has been supported by the discovery of certain MHC-II specific neoantigens in tumor cells and their necessities in promoting successful anti-tumor response [48]. Considering the negative regulation of MHC-II by PLK1, our study further supports this notion in LUAD. Despite that, the mechanistic part behind this regulation is missing and requires further exploration.

Considering the predominant immunosuppressive functions of PLK1 in LUAD and other cancer types, treatment focuses on targeting PLK1 may decelerate tumor growth by boosting immune response in TME. However, it is noteworthy that PLK1 may be negatively correlated with PD-L1 in certain situations [20, 21], devaluing the monotherapy with PLK1 inhibitor as this approach will unexpectedly elevate immune checkpoints. Thus, combination treatment with PLKi and immune checkpoints blockade should exhibit premium effect, and this assumption remains to be investigated by future studies.

## Supporting information

Gating strategy for flow cytometry experiments

## Acknowledgement

This study is supported by NIH R01 CA157429 (XL), R01 CA196634 (XL), R01 CA264652 (XL), R01 CA256893 (XL). This study is also supported by the Biospecimen Procurement & Translational Pathology, Biostatistics and Bioinformatics, Redox Metabolism, Flow Cytometry and Immune Monitoring Shared Resources, and OncoGenomics Shared Resource Facility of the University of Kentucky Markey Cancer Center (P30CA177558).

## Data availability

All data are included in this article. Bulky RNA-seq data was deposited in GEO under accession number GSE206644.

---

**Figure S1.**
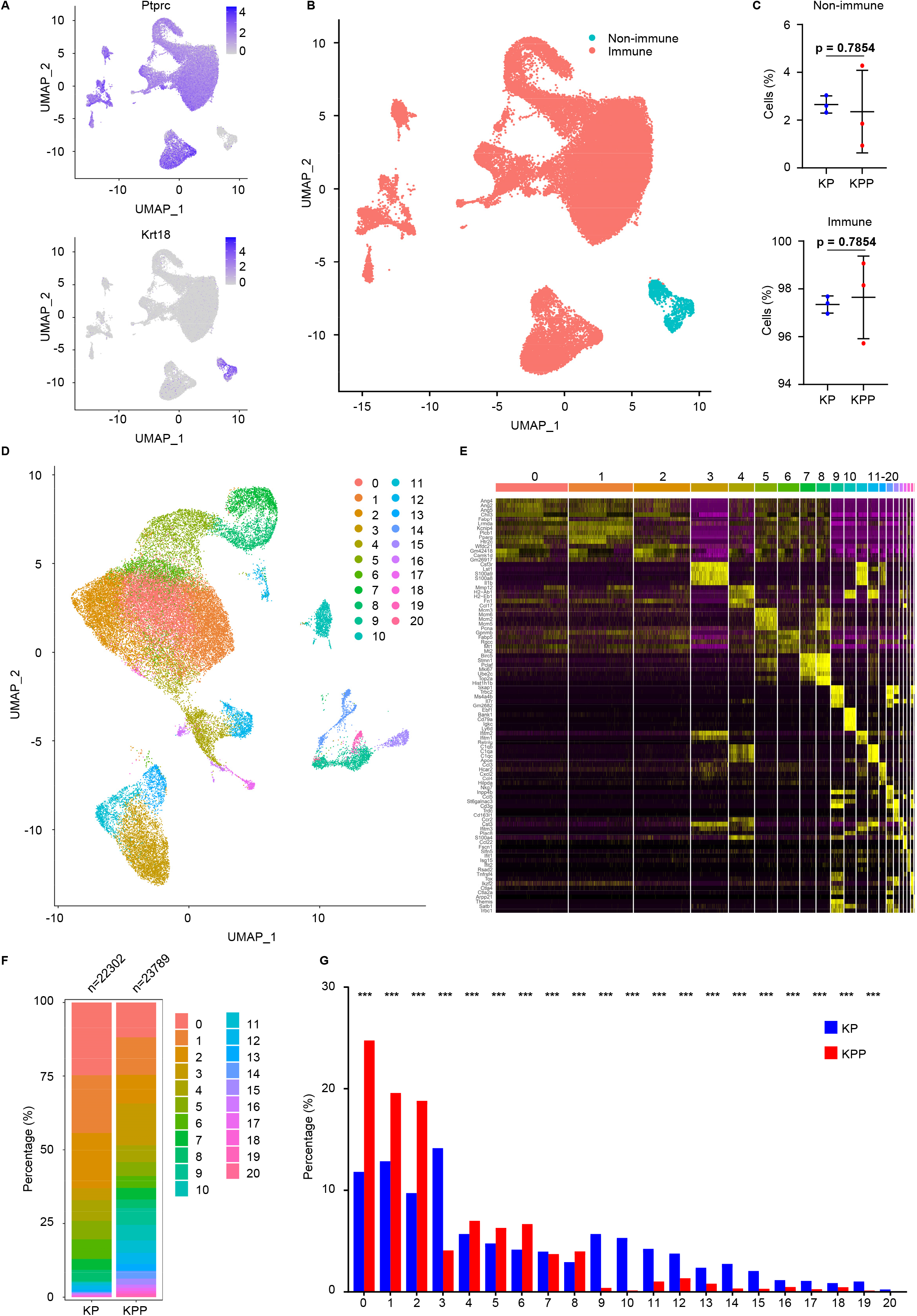
Single-cell RNA-seq analysis. **A,** Representative genes and functional markers used for classification of immune cells (Ptprc) and non-immune cells (Krt18). **B,** UMAP of immune and non-immune cells in KP and KPP. **C,** Comparison of immune and non-immune cells proportions between KP and KPP (n=3). **D,** UMAP of immune cells undergoing unsupervised clustering in KP and KPP. **E,** Heatmap of top 5 signature genes of each cluster in D. **F,** Proportions of unsupervised clusters in KP and KPP. **G,** Comparison of unsupervised clusters proportions between KP and KPP. Statistical method: two-sample binomial test. ***, FDR < 0.001.

**Figure S2.**
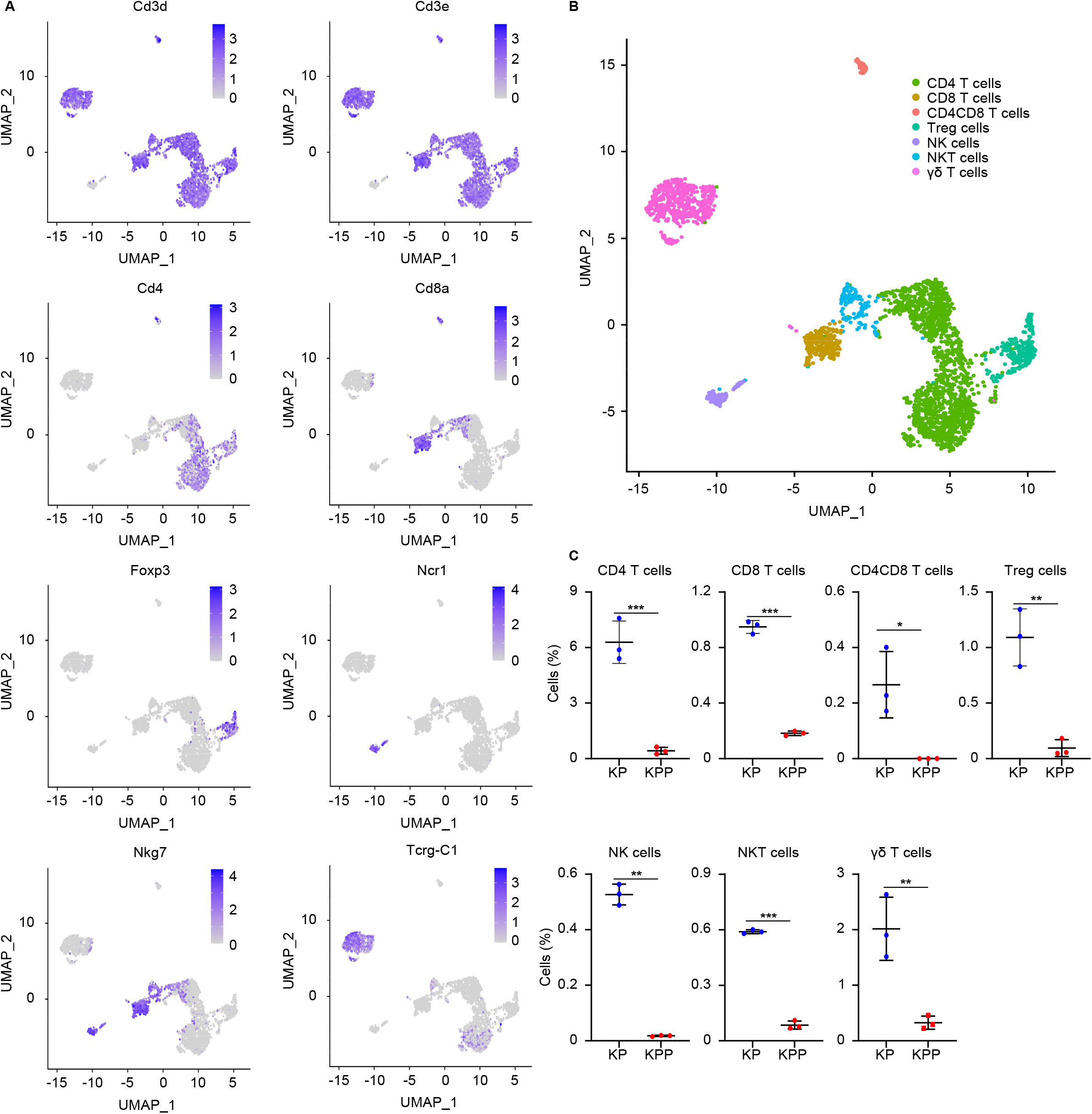
Single-cell RNA-seq analysis of T cell and NK cell populations. **A,** Representative genes used for identifying T cell and NK cell populations. **B,** UMAP projection of T cell and NK cell populations. **C,** Comparison of proportions (in total immune cells) of T cell and NK cell populations denoted in B between KP and KPP (n=3). *, p < 0.05. **, p < 0.01. ***, p < 0.001.

**Figure S3.**
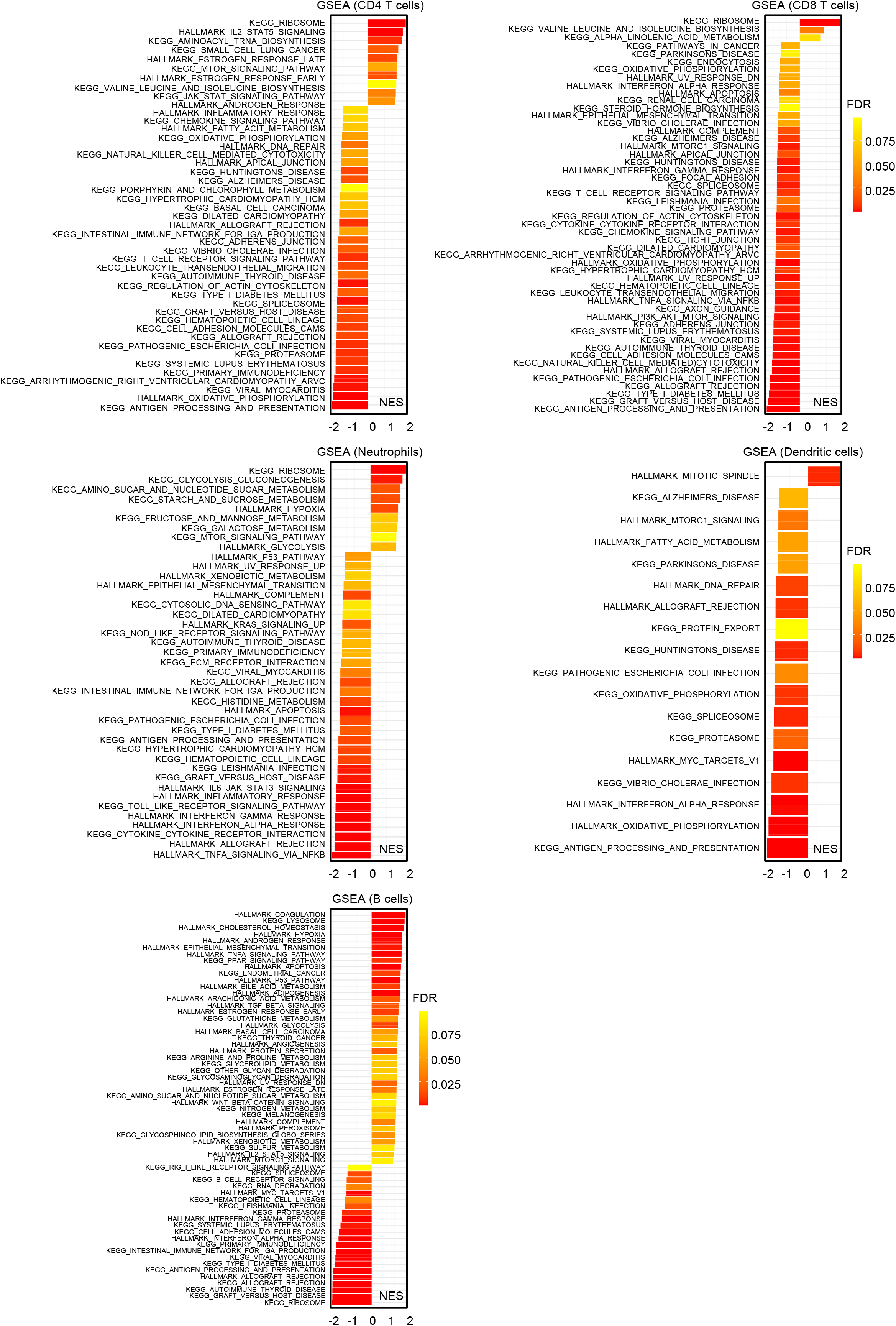
GSEA results of immune cell populations. Significant pathways (KPP vs KP, FDR < 0.1) in the indicated immune cell populations are shown.

**Figure S4.**
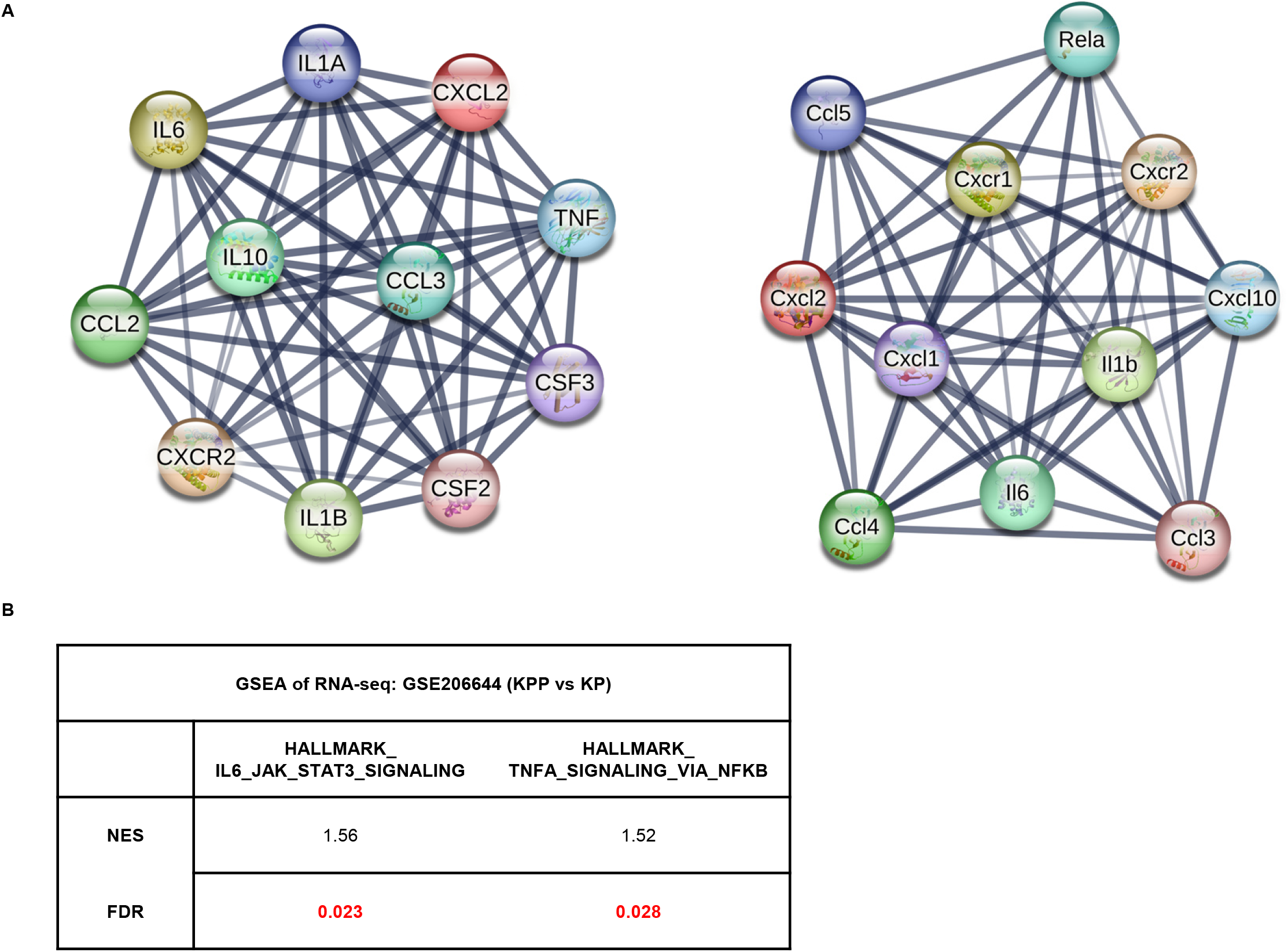
RNA-seq analysis of KP and KPP cells. **A,** STRING analysis of functional genes associated with CXCL2 in human and mice. **B,** GSEA table of two potential pathways regulating CXCL2 secretion in KPP cells, identified by our previously published RNA-seq data (GSE206644, KPP vs KP). FDR < 0.05 indicates a significance.

**Figure S5.**
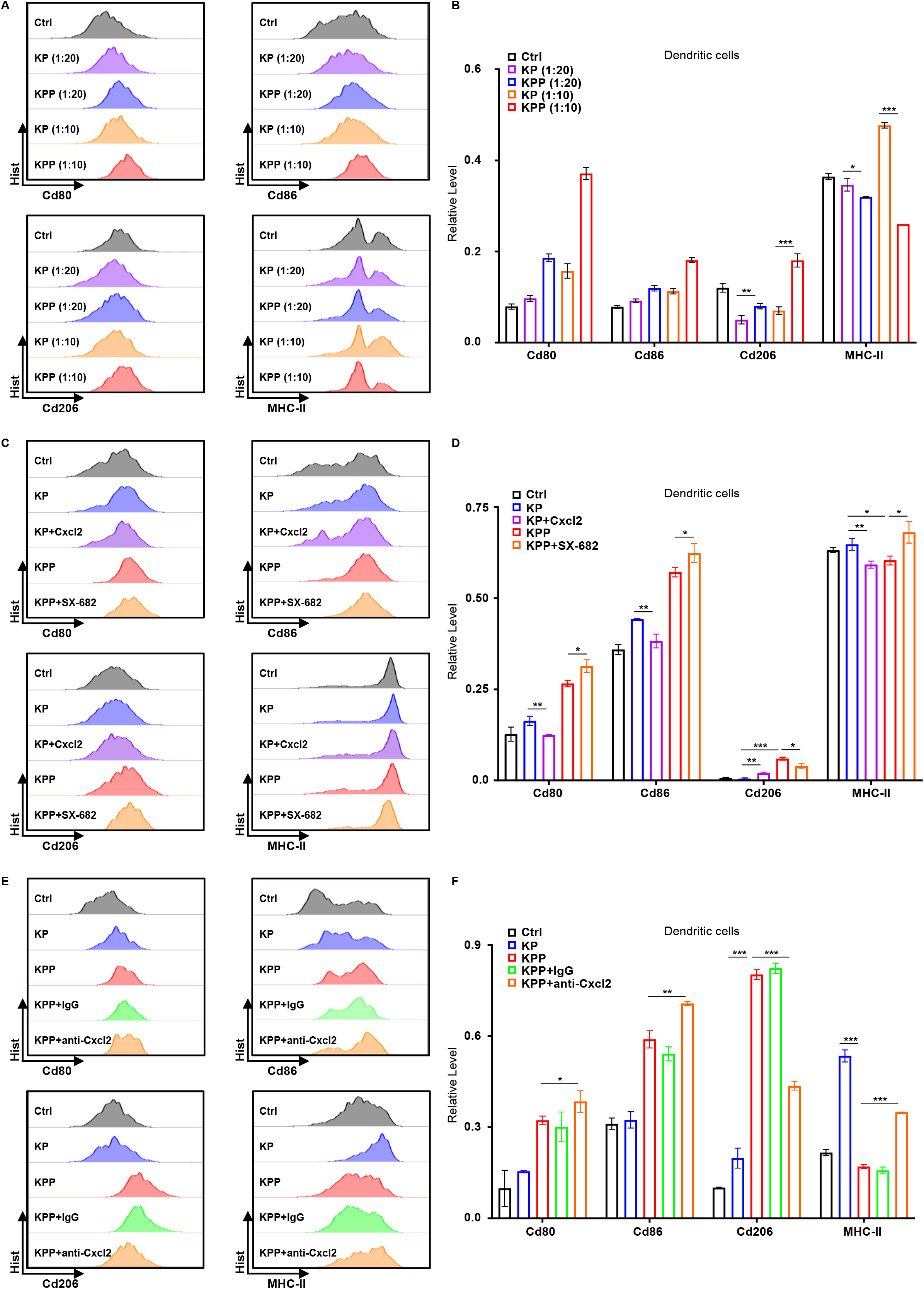
Coculture experiments of dendritic cells. **A,** Flow cytometry analysis of indicated genes in dendritic cells cocultured with KP or KPP at the indicated ratio for 48 hours. **B,** Quantification of A (n=3). **C,** Flow cytometry analysis of dendritic cells cocultured with conditioned medium from KP or KPP for 48 hours, with or without recombinant Cxcl2 (2ug/ml) or CXCR1/CXCR2 inhibitor SX-682 (1µM) treatment. **D,** Quantification of C (n=3). **E,** Flow cytometry analysis of dendritic cells cocultured with conditioned medium from KP or KPP for 48 hours, with or without anti-Cxcl2 neutralization antibodies (5ug/ml) treatment. **F,** Quantification of E (n=3). *, p < 0.05. **, p < 0.01. ***, p < 0.001.

**Figure S6.**
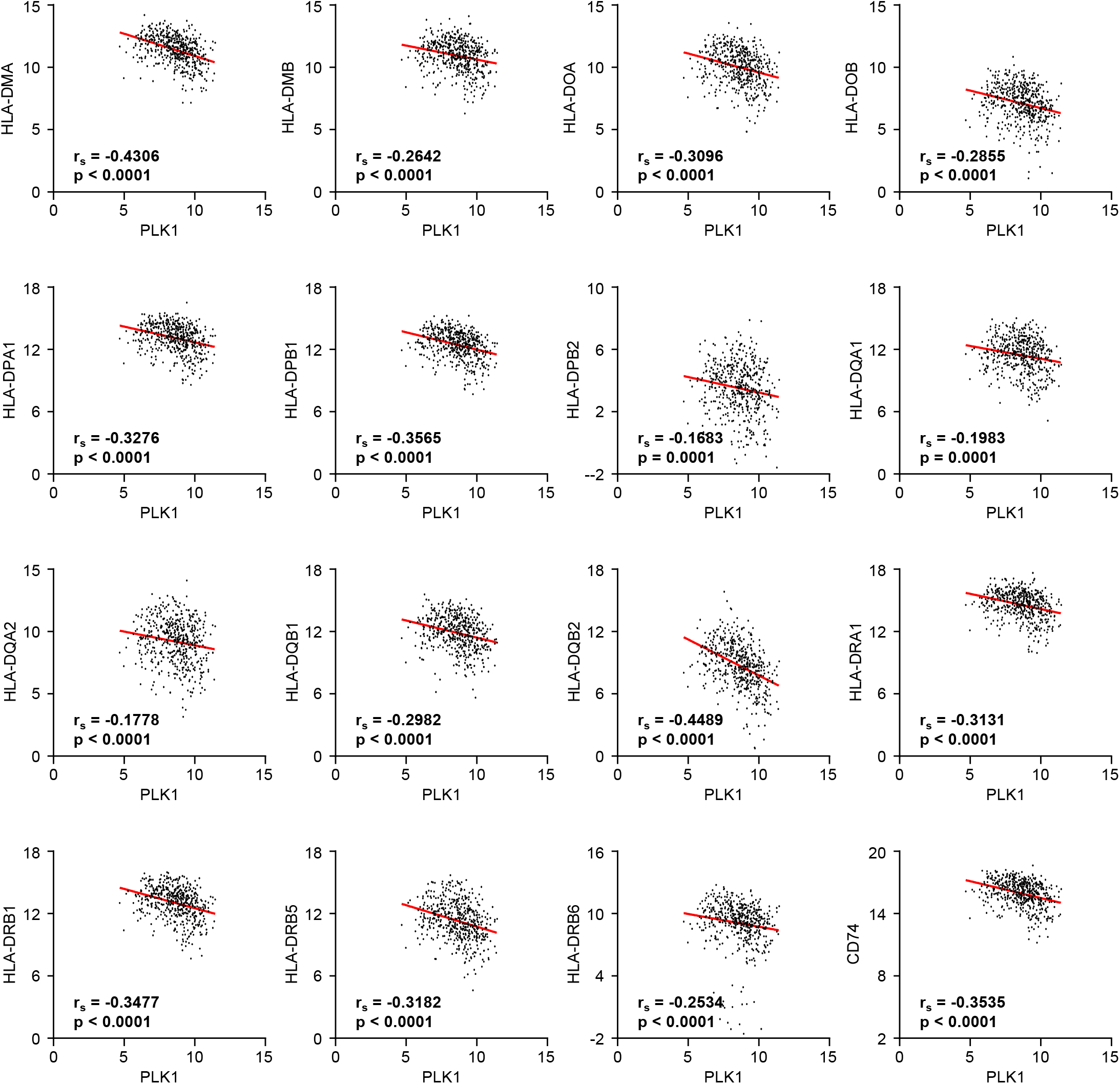
Correlation analysis of PLK1 and MHC-II genes. Patients’ data (Log_2_^RSEM^) are collected from TCGA-LUAD. r_s_, spearman coefficient.

