## Supplementary figures and images for "Single-cell analysis characterizes PLK1 as a catalyst of an immunosuppressive tumor microenvironment in LUAD"

### Gating strategy for flow cytometry experiments

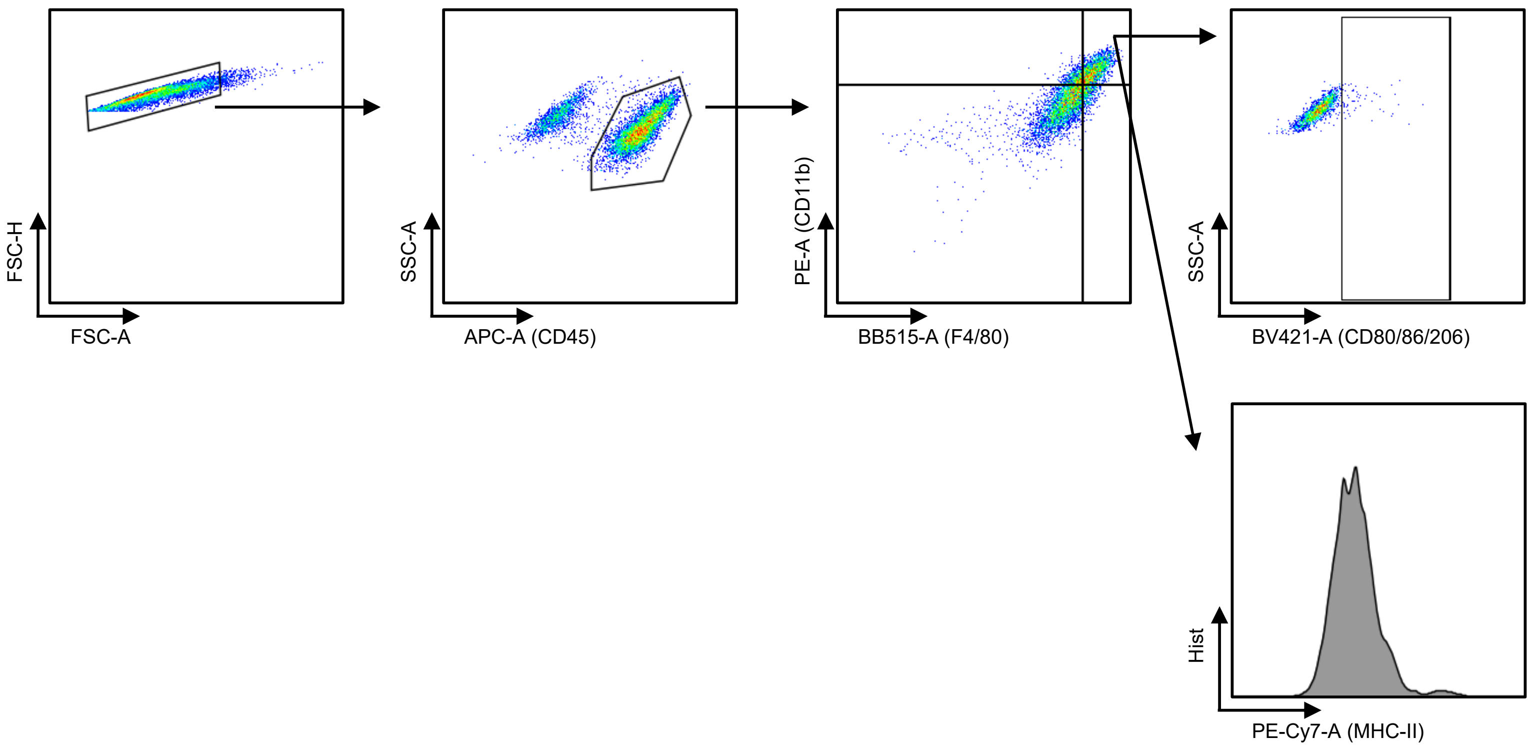

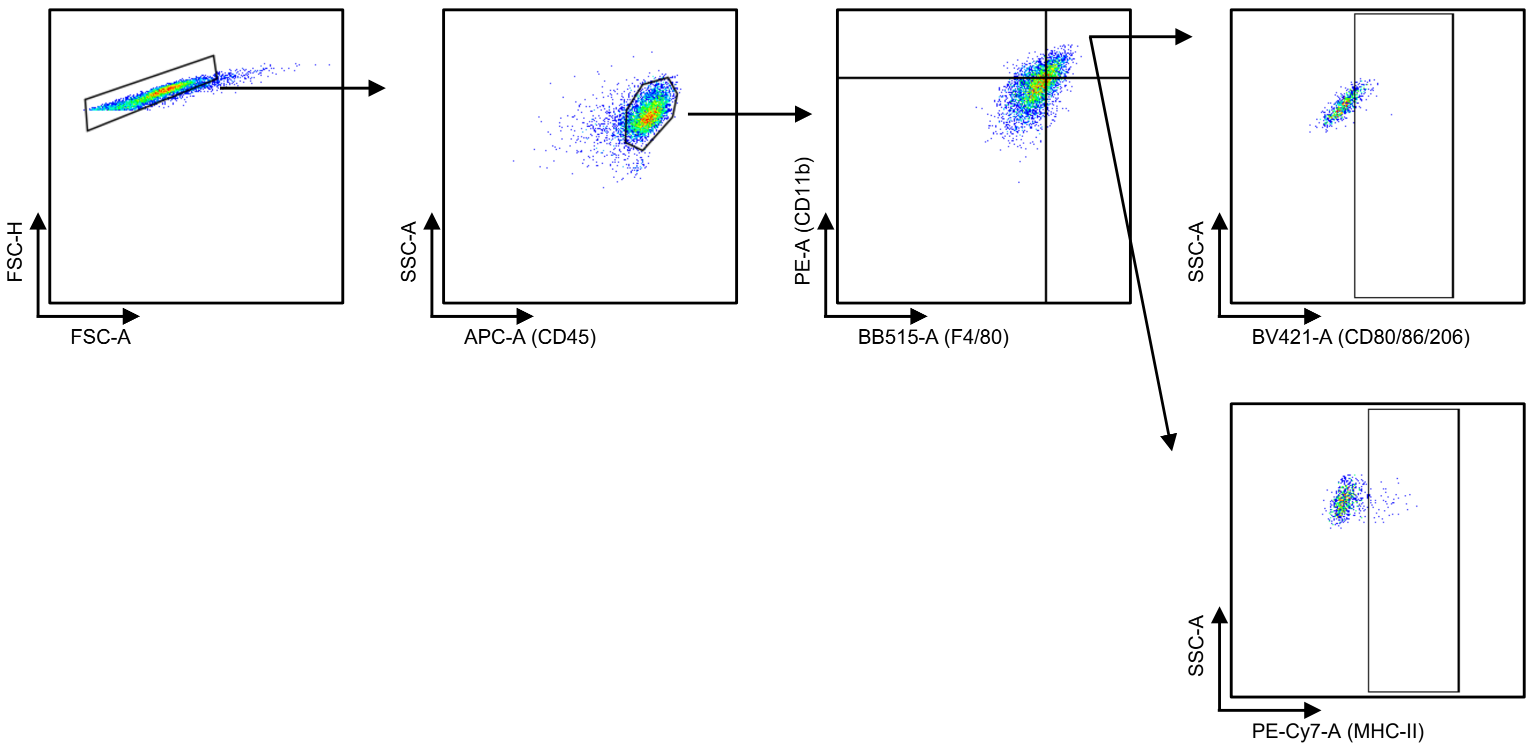

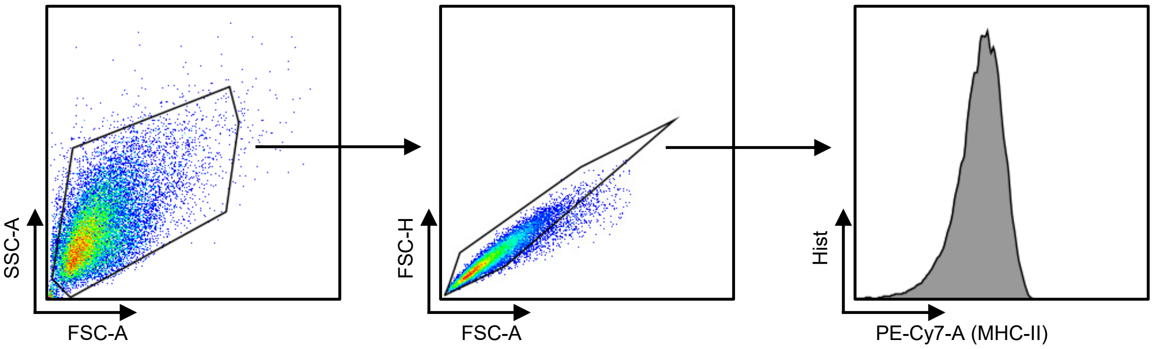

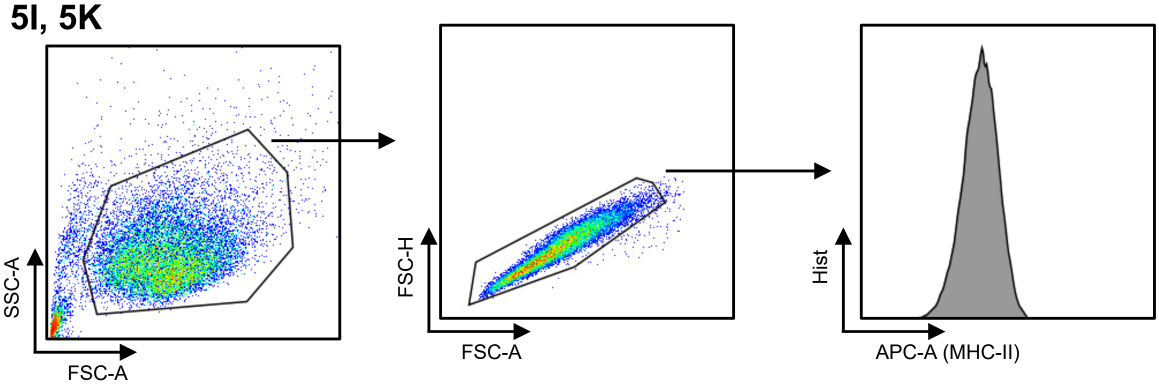

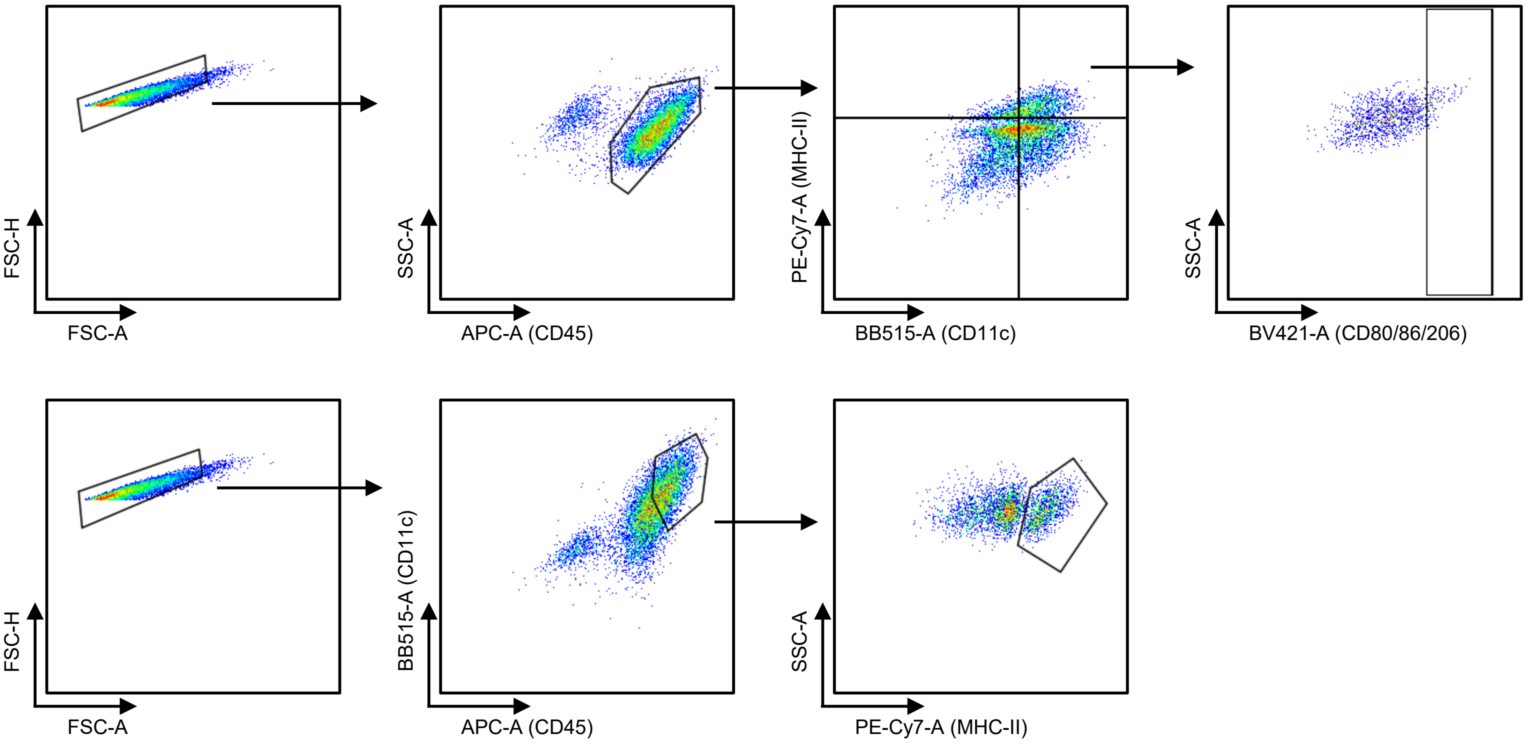

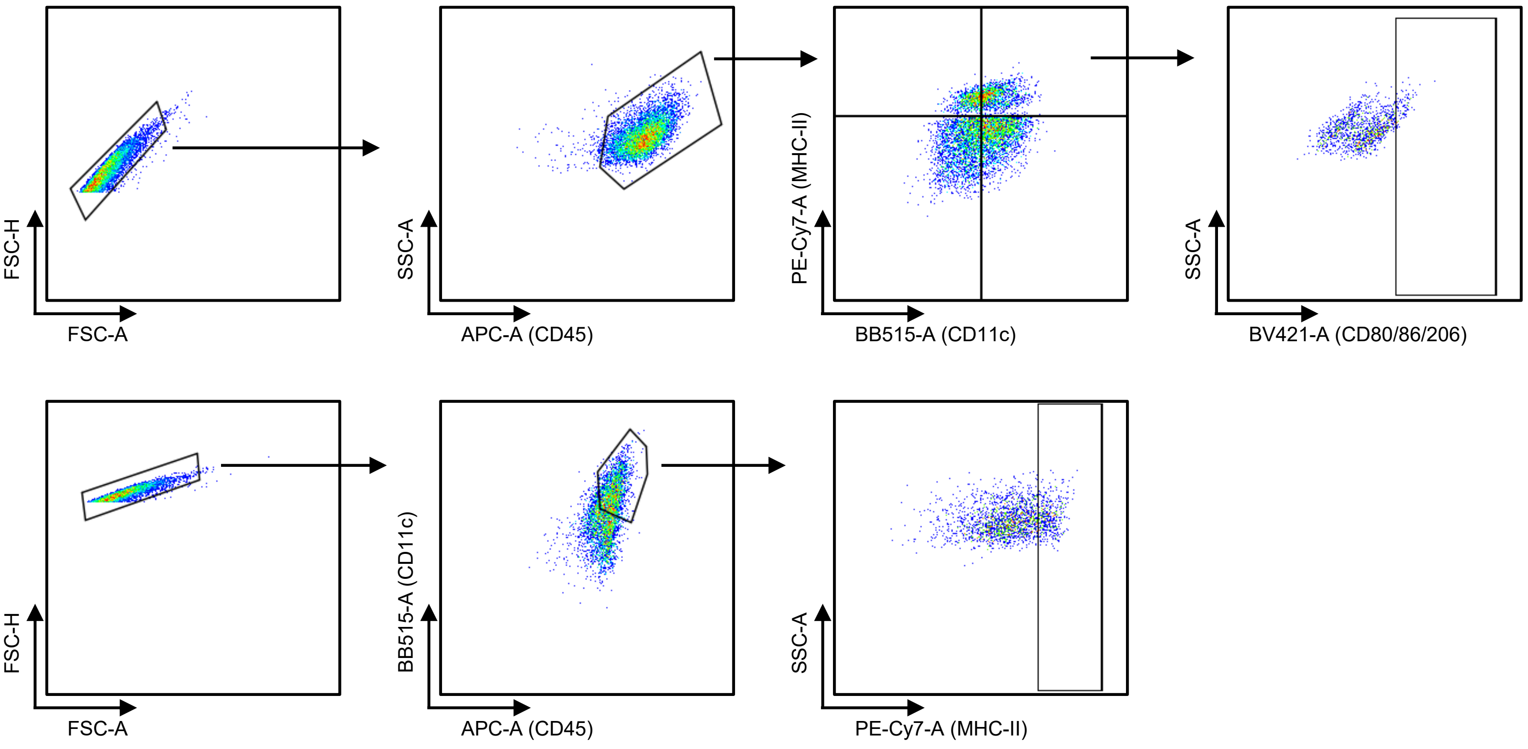
